# Chitosan nanocapsules with *Alstonia boonei* extract modulate the immune system in Wistar rats

**DOI:** 10.1101/2024.10.30.621048

**Authors:** Fonye Nyuyfoni Gildas, Eya’ane Meva Francois, Ninpa Kuissi Chris Rosaire, Bamal Hans-Denis, Fannang Simone Veronique, Beglau Thi Hai Yen, N. A. Fetzer Marcus, Ntoumba Agnes Antoinette, Nguemfo Edwige Laure, Hzounda Fokou Jean Batiste, Djuidje Annie Guilaine, Kuinze Kojap Augustine, Tako Djimefo Alex Kevin, Tchangou Njiemou Armel Florian, Evouna Danielle Ines Madeleine, Dongmo Alain Bertrand, Paboudam Gbambie Awawou, Janiak Christoph

**Affiliations:** Department of Chemistry, Faculty of Science, University of Yaounde I, PO Box 819 Yaounde, Cameroon; Department of Pharmaceutical Sciences, Faculty of Medicine and Pharmaceutical Sciences, University of Douala, PO Box 2701, Douala, Cameroon; Institute for Inorganic and Structural Chemistry, Heinrich Heine University Düsseldorf, 40204 Düsseldorf, Germany; Department of Animal Biology and Physiology, Faculty of Science, University of Douala, PO Box 24157, Douala, Cameroon; Department of Biological Sciences, Faculty of Medicine and Pharmaceutical Sciences, University of Douala, Cameroon, Cameroon; Clinical Biology, Faculty of Medicine and Pharmaceutical Sciences, University of Douala, PO Box 2701 Douala, Cameroon

**Keywords:** *Alstonia boonei*, Nanocapsule, Acute toxicity, Immunomodulatory activity

## Abstract

The development of biosynthetic methods for nanoparticles using plants presents an exciting opportunity to better utilize our rich and diverse medicinal flora. We are particularly interested in creating nanocapsules using *Alstonia boonei*, a Cameroonian plant known for its immunomodulatory properties.

Chitosan nanocapsules were synthesized from the methanol/dichloromethane extract of the powdered stem bark of *A. boonei* after harvest and drying. The encapsulation of the secondary metabolites was achieved using the ionic gelation method, which involved the agitation of a chitosan solution, extract, and tripolyphosphate. Subsequently, the encapsulation efficiency was calculated. Infrared spectroscopy identified the various functional groups present in the nanocapsules. The acute toxicological profile of these chitosan nanocapsules at a limit dose of 2000 mg/kg, along with their immunomodulatory activities, was evaluated in Wistar rats. The immunomodulatory potential was assessed in dexamethasone-induced immunosuppressed rats by measuring total blood count, delayed-type hypersensitivity response, and hemagglutinating antibody titre between groups of animals after 14 days of treatment.

The data collected on the synthesis and characterization confirmed the formation of nanocapsules. This was evidenced by infrared spectroscopy and an entrapment efficiency of 69%. Powder X-ray diffraction confirmed the presence of chitosan in the polymer material. SEM imaging further confirmed the formation of nanocapsules. The toxicological profile of these nanocapsules was found to be satisfactory. Administration of chitosan nanocapsules containing *Al. boonei* methanol/dichloromethane extracts at doses of 100 mg/kg, 200 mg/kg, and 500 mg/kg body weight significantly prevented dexamethasone-induced immunosuppression in rats. This was achieved by increasing the parameters of total blood count (hematocrit, mean corpuscular volume, platelets, lymphocytes, and granulocyte counts), hemagglutinating antibody titre values, and delayed type hypersensitivity response induced by chicken red blood cells. However, doses of 500 mg/kg of crude *A. boonei* extract and 500 mg/kg body weight of empty chitosan nanocapsules did not show this effect.

The nanocapsules generated from the extracts of chitosan and *A. boonei* are responsible for immunostimulatory activity and possess therapeutic potentials for the prevention of depressed immune depressed conditions with satisfactory safety at acute dose.

## 1. Introduction

Innate immunity consists of a series of host defenses that provide the initial response to pathogens or injuries. These responses are phylogenetically ancient and have evolved to manage pathogens that are frequently encountered but rarely cause diseases. Immunomodulatory drugs alter the immune system response by either enhancing or suppressing serum antibodies, antigen recognition and phagocytosis, lymphocyte proliferation, antigen-antibody interactions, mediator release due to immune response, and modification of target tissue responses [1]. The immune system plays a central role in many chronic diseases. Understanding altered immune function in chronic diseases such as cancer, rheumatoid arthritis, inflammatory bowel disease, asthma, multiple sclerosis, diabetes, heart disease, and others has not only elucidated the mechanisms underlying these diseases, but also suggested new therapies that can positively impact patients by reducing morbidity and mortality [2].

In Africa, thousands of plants are commonly used to treat diseases, and their role in the treatment and prevention through the strengthening of the immune system is of significant interest [3]. The *Apocynaceae* family, based on chemical studies, has been shown to possess extremely rich metabolites that could play a crucial role in drug discovery by providing molecules with potential as templates for therapeutic drug development [4]. Pharmacological studies on these plants have demonstrated various biological activities, including antioxidant, anti-inflammatory, antimicrobial, antimalarial, analgesic, hypotensive, antidiabetic, and anti-parasitic properties, making them useful to prevent or cure various pathologies [5].

Recent advancements in the interdisciplinary field of nanotechnology have spurred innovative approaches in the design of nano-sized drug carriers and delivery systems. This progress has led to the creation of new nanometric materials applicable in biology, biotechnology, medicine, and medical technology [6]. A notable example is the synthesis of nanocapsules from chitosan, a polymer derived from the exoskeletons of crustaceans. Chitosan is gaining significant attention due to its abundance, biocompatibility, biodegradability, low toxicity, anti-inflammatory properties, ability to improve penetration, efficacy in drug delivery and targeting, wound healing, and immunoenhancing capabilities [7].

*Alstonia boonei* (De Wild), a member of the Apocynaceae family, is locally known as kokmot or njie in Bassa‘a and ekouk or nfoul in Ewondo. This plant, part of the rich Cameroonian flora, is valued for its anti-inflammatory, antiparasitic, and antimicrobial properties, which are probably due to its immune-stimulant potential [2]. This study aims to alleviate immunomodulatory-mediated diseases by combining chitosan with *A. boonei* extract in nanocapsules.

## 2. Materials and methods

### Plant Material and Extraction

The stem bark of the plant was harvested in Mbalmayo, Center region, Cameroon and identified as *Alstonia boonei* at the National Herbarium of Cameroon in Yaoundé, with reference number 43364/HNC. The plant material was dried in the shade, away from sunlight, for 2-3 weeks and then ground to a fine powder. A 100 g sample of the powder was subjected to double maceration, each cycle lasting 48 hours, in 1.5 liters of a methanol/dichloromethane (CH_3_OH/CH_2_Cl_2_; 70/30) mixture.

The filtrate was concentrated at 40 °C under reduced pressure using a rotary evaporator and then dried *in vacuo* [8]. The extraction yield was calculated using formula 1:

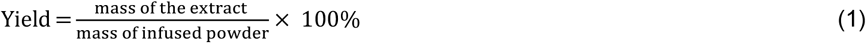

#### Qualitative phytochemical analysis of the extract

The extract of *A. boonei*’s trunk bark was assessed for the presence of different primary and secondary metabolites groups and classes, namely reducing sugars, phenolic compounds including flavonoids and coumarins, alkaloids, triterpenes including saponins and steroids, following le Thi et al. protocols [9].

#### Synthesis of Chitosan Nanocapsules

Two chitosan solutions were prepared at a concentration of 10 mg/mL by adding 50 g of chitosan powder to 5 liters of acetic acid at 2% (v/v). The pH was adjusted within the range 4.5 - 5.5. The methanol/dichloromethane extract of *A. boonei* (20 mg) was added to a chitosan solution. Each solution was then mixed with an equal volume of a 4 mg/mL sodium tripolyphosphate solution using a magnetic stirrer at 7200 rpm. Upon the formation of nanocapsules (indicated by the opacification of the mixture), the suspensions were subjected to sonication at a frequency of 120 Hz and an amplitude of 20% for 5 minutes, followed by centrifugation at 13000 rpm [10].

#### Encapsulation efficiency of extract-containing chitosan nanocapsules

The encapsulation efficiency of chitosan nanocapsules (CNC) containing *A. boonei* extract was evaluated by determining the total phenol content using the Folin-Ciocalteu method, as previously described [11]. Briefly, 1 mL of the CNC solution was added to a 10% Folin-Ciocalteu solution and incubated for 30 minutes. Subsequently, 4 mL of 0.7 M sodium bicarbonate solution was added, and the mixture was agitated and stored at room temperature for 2 hours. The absorbance of the mixture was measured at 765 nm using UV-Vis spectrophotometry, which allowed the calculation of phenol concentrations in milligram equivalents of gallic acid per gram of the methanol/dichloromethane extract (mg EQ/gE) [11]. The encapsulation efficiency was then calculated using formula 2:

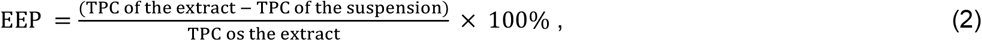

With

EEP: Efficiency of polyphenol encapsulation in %

TPC: Total phenolic content

## Methods

Ultraviolet-Visible Spectrophotometry was monitored using an aliquot of 2 mL of bio composite suspension on a UV-Vis GENESYS 10S UV–Vis Spectrophotometer (Thermo). Measurements were made between 200 and 800 nm.

Fourier Transform Infrared spectroscopy was performed using a Bruker Tensor 37 with attenuated total reflection, ATR unit by scanning between 600 and 4000 cm^−1^.

Powder X-Ray Diffraction measurements of chitosan-*A. boonei* nanocomposites were carried out using a Bruker D2 Phaser powder diffractometer (Cu K-Alpha1 [Å] 1.54060, K-Alpha2 [Å] 1.54443, K-Beta [Å] 1.39225) by preparing a thin film on a low background silicium sample holder (Supplement 1).

Scanning Electron Microscopy (SEM) investigations were performed with a Jeol JSM - 6510LV QSEM Advanced electron microscope with a LaB6 cathode at 20 kV

### Animal Material

The study was carried out on young adults (8 to 12 weeks old) Wistar rats (*Rattus norvegicus* weighing between 120 and 180 grams), housed in the animal facility of the Faculty of Medicine and Pharmaceutical Sciences of the University of Douala. Rats were divided into batches and left for acclimatization prior to testing. Ethical clearance (Nr. 3224 CEI-Udo/06/2022/T) was obtained from the Institutional Research Ethics Committee for Human Health of the University of Douala (CEI-UDo).

### Acute toxicity study

The acute toxicity study was carried out according to the guidelines of the Organization for Economic Cooperation and Development (OECD) protocol directives N°423, with slight modifications as described elsewhere [12]. Briefly, two batches of female rats (n=3) were used. The rats fasted for 12 h prior to the test. The test batch received 2000 mg/kg body weight (b. w.) of CNC while the control batch received distilled water (10 mL/kg b.w.). Clinical parameters were observed with particular attention during the first 30 minutes, then the fourth and eighth hours after administration. Subsequent observations were made every 24 h, regarding the modification of the skin, fur, mucus, behavior, lethargy, sleepiness, and coma. The rats were also weighed every 48h (2 days). On day 14, the rats were sacrificed under anesthesia with an ether solution. Vital organs, namely the heart; liver, kidneys, lung, and spleen, were removed and their relative masses were calculated against the body weight.

### In vivo assessment of the immunomodulatory activity of chitosan nanocapsules

In vivo assessment of the immunomodulatory activity was assessed according to Ukpo et al., protocol with few modifications. Well-fed Wistar rats aged 8 - 12 weeks and weighing between 150 and 200 g were divided into 8 batches named B0 to B7. B0 served as a sham control and B1 was the negative control batch and was treated with distilled water. Immunosuppression was induced in B1-7 by intraperitoneal injection of dexamethasone (5 mg/kg b.w.), twice daily, for 3 days. On day 4, blood samples (2 mL) were collected from rats to confirm immunosuppression, then immunosuppressed rats were treated with different substances: B2 received levamisole (50 mg/kg b.w.) and served as positive control, while B3 was treated with empty chitosan nanocapsules at 500 mg/kg and B4 was treated with *A. boonei* crude methanol/dichloromethane extract at 500 mg/kg b.w. Batches 5, 6 and 7 were treated with increasing doses of CNC containing the methanol/dichloromethane extract: 100, 200 and 500 mg/kg b.w., respectively. Blood samples (2 mL) were collected, and hematological parameters were assessed using a hematology analyzer for complete and differential blood cell counts [13].

### Determination of delayed-type hypersensitivity responses (DHT)

DHT was determined according to Wardani and Sudjarwo reports. On day 7 of the study, rats in test groups were primed by subcutaneously injecting 0.1 mL of suspension containing 1×10^8^ red blood cells obtained from chicken sacrificed in a local slaughterhouse, into the right hind footpad. The contralateral paw also received an equal volume of 0.1% phosphate buffered saline (PBS). The administration of methanol/dichloromethane extracts and the drug continued until day 14. The animals were then challenged by subcutaneous injection of 0.1 mL of 1×10^8^ chicken red blood cells (CRBCs) into the left hind footpad. The extent of the DTH response was assessed by measuring the thickness of the footpad at 4-, 8-, and 24-hours post-challenge using a digital vernier caliper. The difference in the thickness of the right and left hind paws was used as a measure of the DTH reaction and expressed as a mean percent increase in thickness/edema [14].

### Hemagglutination antibody titre test

Rats in the test groups were immunized with an I.P. injection of 0.5 mL of CRBCs on day 7 of the experiment. The administration of drugs and methanol/dichloromethane extracts was carried out for another 7 days until day 14 and blood samples were collected by cardiac puncture. Blood was centrifuged at 1609.92 x*g* to obtain serum. The antibody titers were then determined using the hemagglutination technique described by Wardani and Sudjarwo, with slight modification [14]. Briefly, serial two-fold serum dilutions were made with normal saline in microtiter plates of 96 - well capacity and CRBCs (25 μL of 1% CRBCs prepared in normal saline) added to each of these dilutions. The hemagglutination plates were then incubated at 37 °C for 1 h and then examined for hemagglutination. The reciprocal of the highest dilution of test serum giving agglutination was taken as the hemagglutination antibody titre (HA units/μL) [14].

## Data Analysis

The data from all tests were recorded using MS Excel 2013 and analyzed using GraphPad Prism software. All data were expressed as means ± SEM values. The ANOVA test was used to test differences between groups. Duncan’s multiple range test was used to analyze differences between mean values and differences were considered statistically significant at p < 0.05.

## Results

### Extraction Yield

Extraction by double maceration in a methanol/dichloromethane (70/30) solvent system was carried out on 500 g of dry *A. boonei* powder, for 48 h. The extraction yielded 30.55 g (6.11% m/m) of a viscous brown substance (Table 1).

**Table 1.**
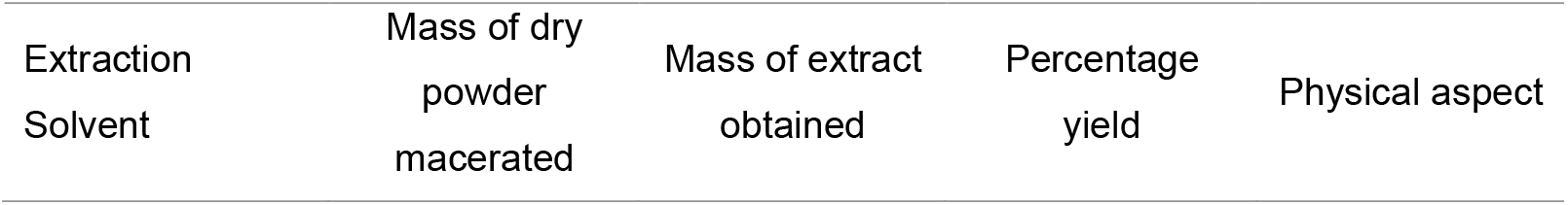
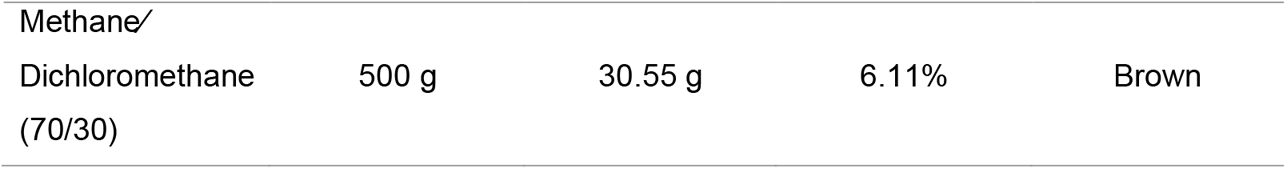
Quantitative outcome from the Methane/Dichloromethane extraction of *Alstonia boonei*.

### Qualitative phytochemical analysis

Identification of secondary metabolites present in the extract of *A. boonei* showed that all groups of secondary metabolites were present in the extract (Table 2).

**Table 2.** Secondary metabolites content of *Alstonia boonei* extract.

| Test | Observation |
| --- | --- |
| Phenols | + |
| Coumarins | + |
| Anthraquinones | + |
| Flavonoids | + |
| Anthocyanidins | – |
| Triterpenes and Steroids | + |
| Saponins | + |
| Alkaloids | + |
| Reducing sugars | + |

(+) present (-) absent

Phenolic compounds were present in the extract, including coumarins and anthraquinones. Flavonoids were also detected, although anthocyanidins were missing. Terpenoids such as triterpenes and saponins were detected, as well as steroids. General tests for alkaloids and reducing sugars were positive.

### Synthesis and characterization of chitosan nanocapsules

Encapsulated *A. boonei* extracts were obtained with an encapsulation efficiency of 69.34% (Table 3).

**Table 3.**
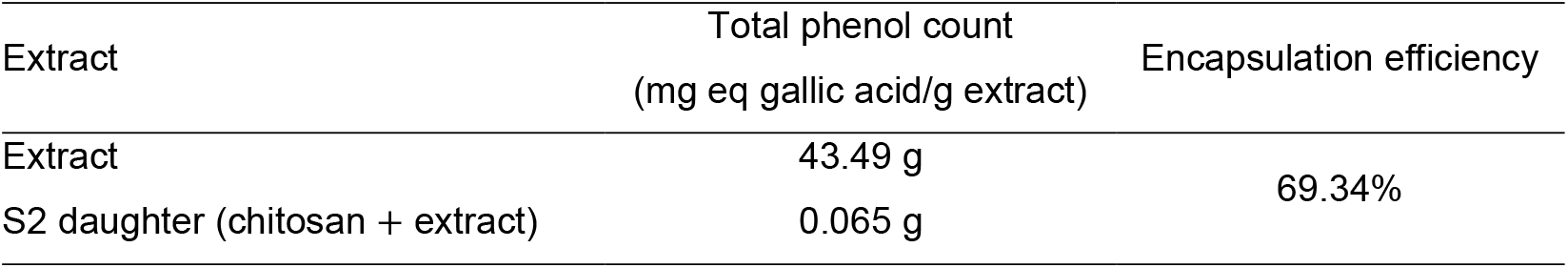
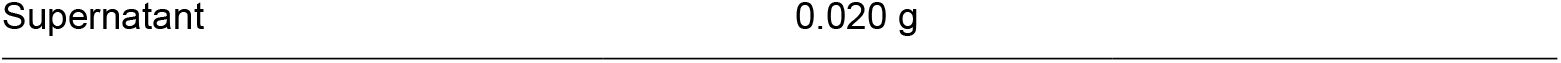
Encapsulation efficiency of CNC.

### Fourier Transform Infrared Spectroscopy analysis

IR analysis was performed on wet samples of chitosan nanocapsules between 4000 cm^-1^ and 450 cm^-1^. A spectrum (Figure 1) was registered showing the different functional groups available at the surface of the chitosan nanocapsules.

**Figure 1.**
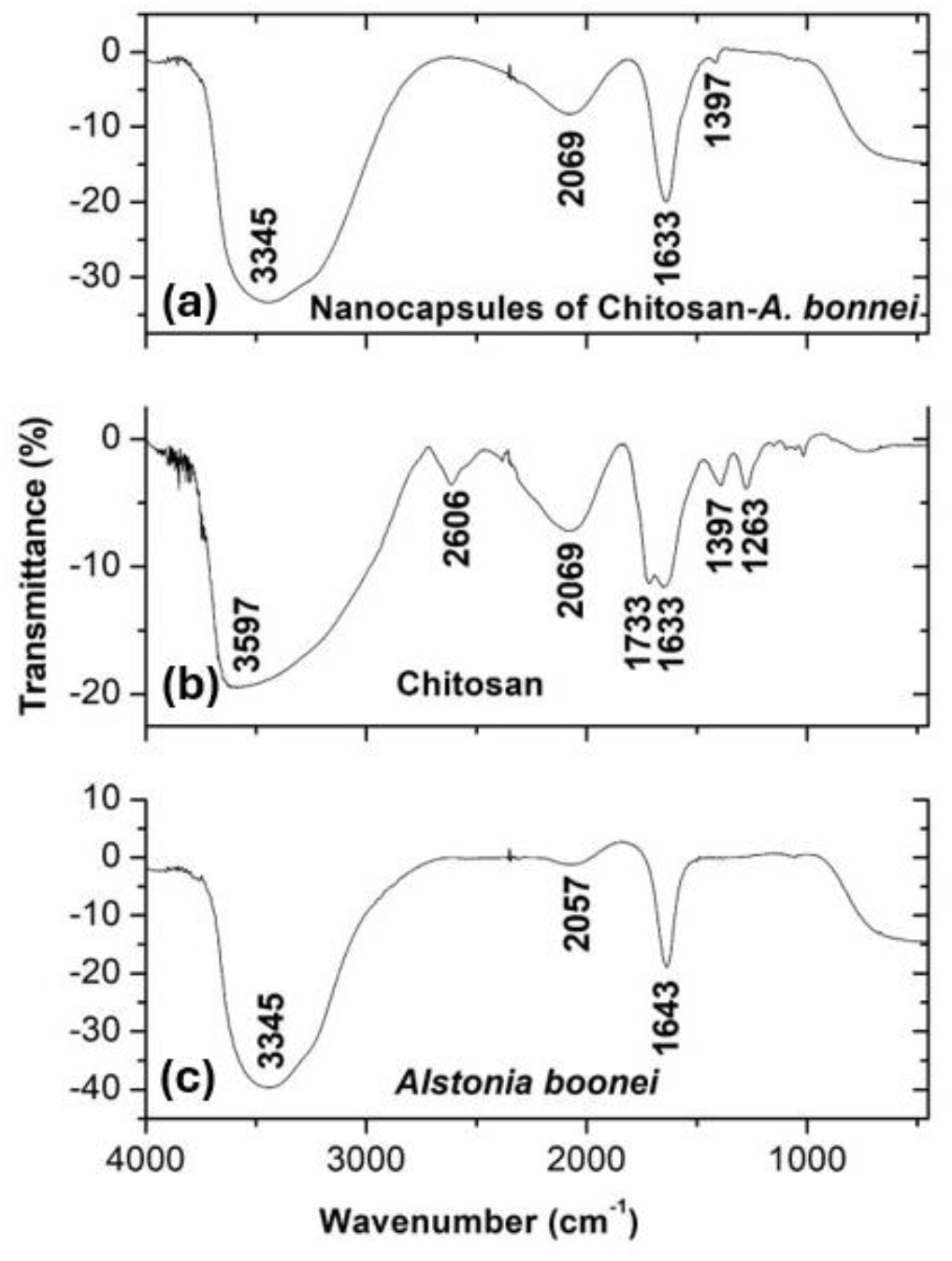
FTIR spectrum of Chitosan-*A. bonnei* nanocapsules (a),, chitosan (b)and *A. bonnie* extract (c)

The IR spectrum of all three substances, namely *A. boonei* extract, chitosan solution and the chitosan nanocapsules, revealed the presence of different chemical moieties in their structures. Similarities were observed at 3345 cm^−1^ for the nanocapsules and *A. boonei* extract, 2069, 1633 and 1397 cm^−1^ for the nanocapsules and chitosan. The broad band at 3300 - 3600 cm^−1^ is indicative of N-H deformation vibrations in amines and O-H stretching vibrations in alcohols and phenols. A very weak peak at 2606 cm^−1^ in the chitosan spectrum corresponds to C-H bond stretching vibrations in alkanes. The spectra of the three solutions also showed a similar weak band at 2069 cm^−1^ for the nanocapsules and chitosan and 2057 cm^−1^ for A. boonei metabolites, characteristic of C-N bonds in amines. A medium band at 1733 cm^−1^ for chitosan is indicative of C=O double bonds in aldehydes and ketones. The bands around 1640 and 1400 cm^−1^ in the substances are characteristic of benzene rings with C=O and C-C bonds. These observations confirm the presence of polyphenols of the extracts and chitosan in the nanocapsules.

The SEM of Chitosan-*A. boonei* nanocapsules image (Figure 2) revealed the formation of encapsulated porous aggregates

**Figure 2.**
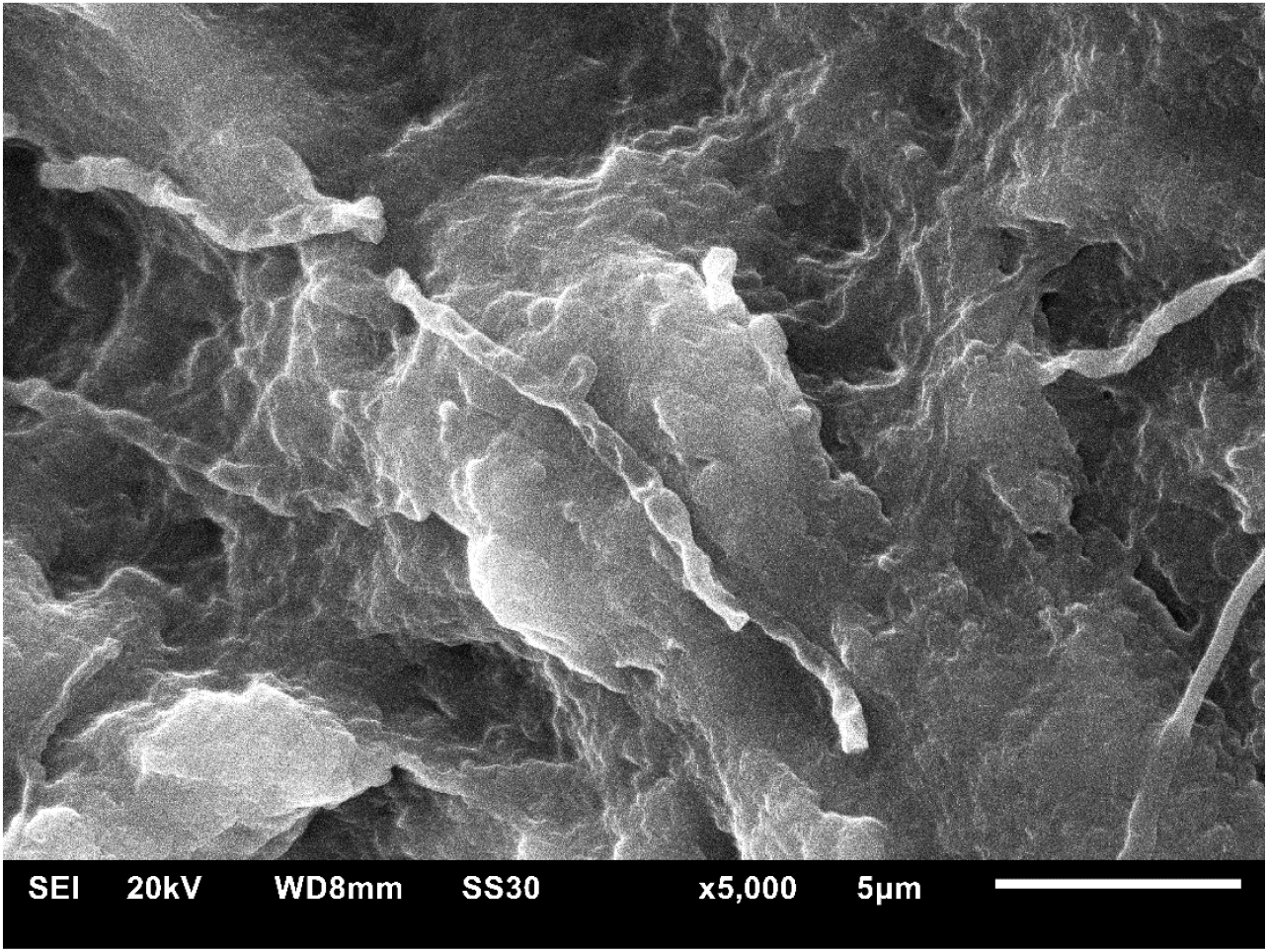
SEM of Chitosan-*A. bonnei* nanocapsules

### Acute toxicological profile of synthesized chitosan nanocapsules

Clinical parameters in rats were monitored after administration of synthesized chitosan nanocapsules at a limit dose of 2000 mg/kg (Table 4). No abnormalities were observed, and no mortality occurred by the 14th day of the evaluation period. This suggests a DL_50_ greater than 2000 mg/kg for CNC with *A. boonei* stem bark extract.

**Table 4.** Assessed clinical parameters in rats.

| Clinical parameters | Negative control | Test batch |
| --- | --- | --- |
| | (Distilled water at $10\text{ mL/kg b.w}$) | (CNC+Extract at $2000\text{ mg/kg}$) |
| Aspect of faeces | N | N |
| Aspect of fur | N | N |
| Diarrhea/vomiting | A | A |
| Respiratory rate | N | N |
| Tremor | A | A |
| Dyspnea/Tachycardia | A | A |
| Agitation | A | A |
| Convulsions/coma/<br>death | A | A |
| DL<sub>50</sub> | - | > 2000 mg/kg |

N : Normal ; A : Absent

The weight variation monitoring of rats subjected to the extract at the limit dose between day 1 and day 14 revealed a normal increase in body weight, with no significant differences between the negative control and the test batch (Figure 3).

**Figure 3.**
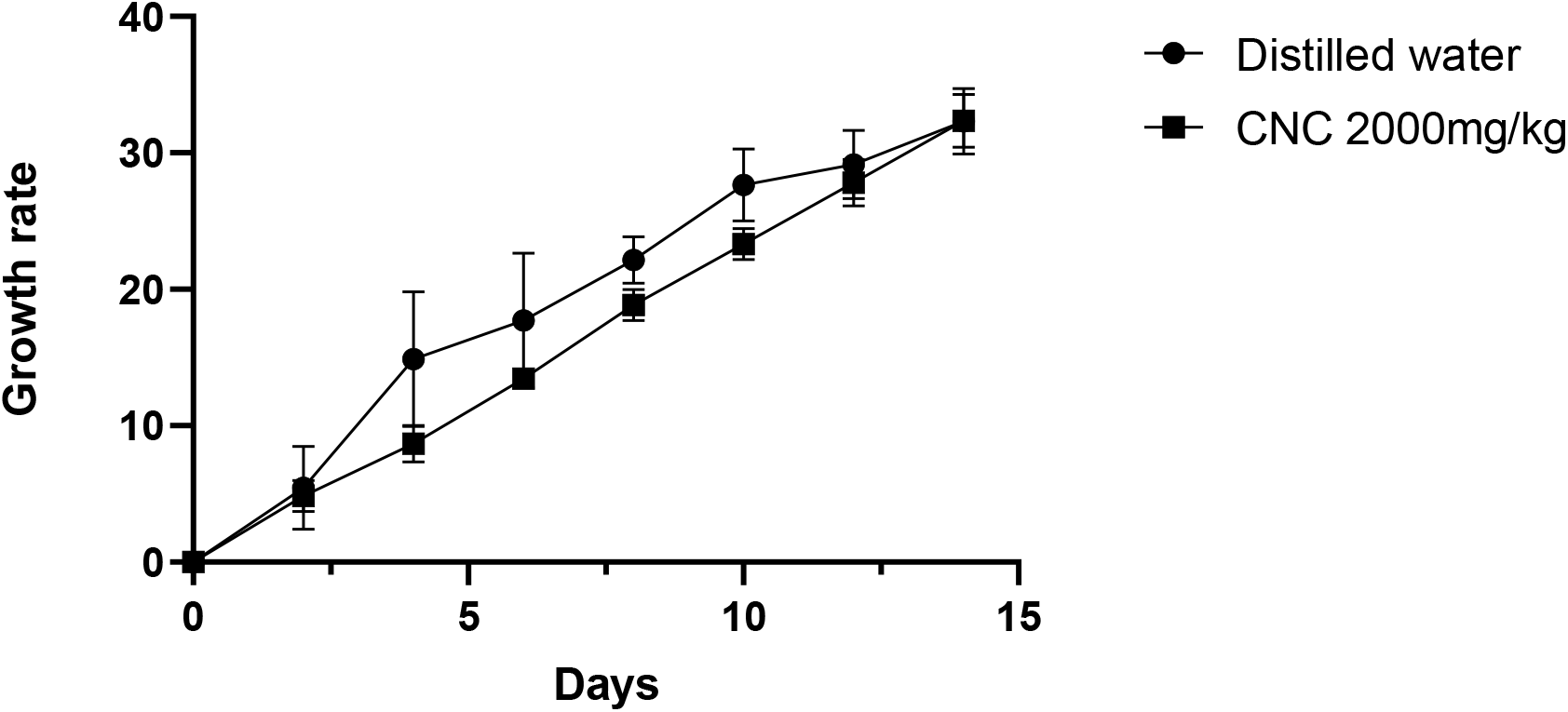
Growth rate of rats

Furthermore, comparative analysis of the relative weight of the internal organs, namely the heart, kidneys, liver, lungs, and spleen, compared to the negative control group revealed no significant differences between the two sets of values (Figure 4).

**Figure 4.**
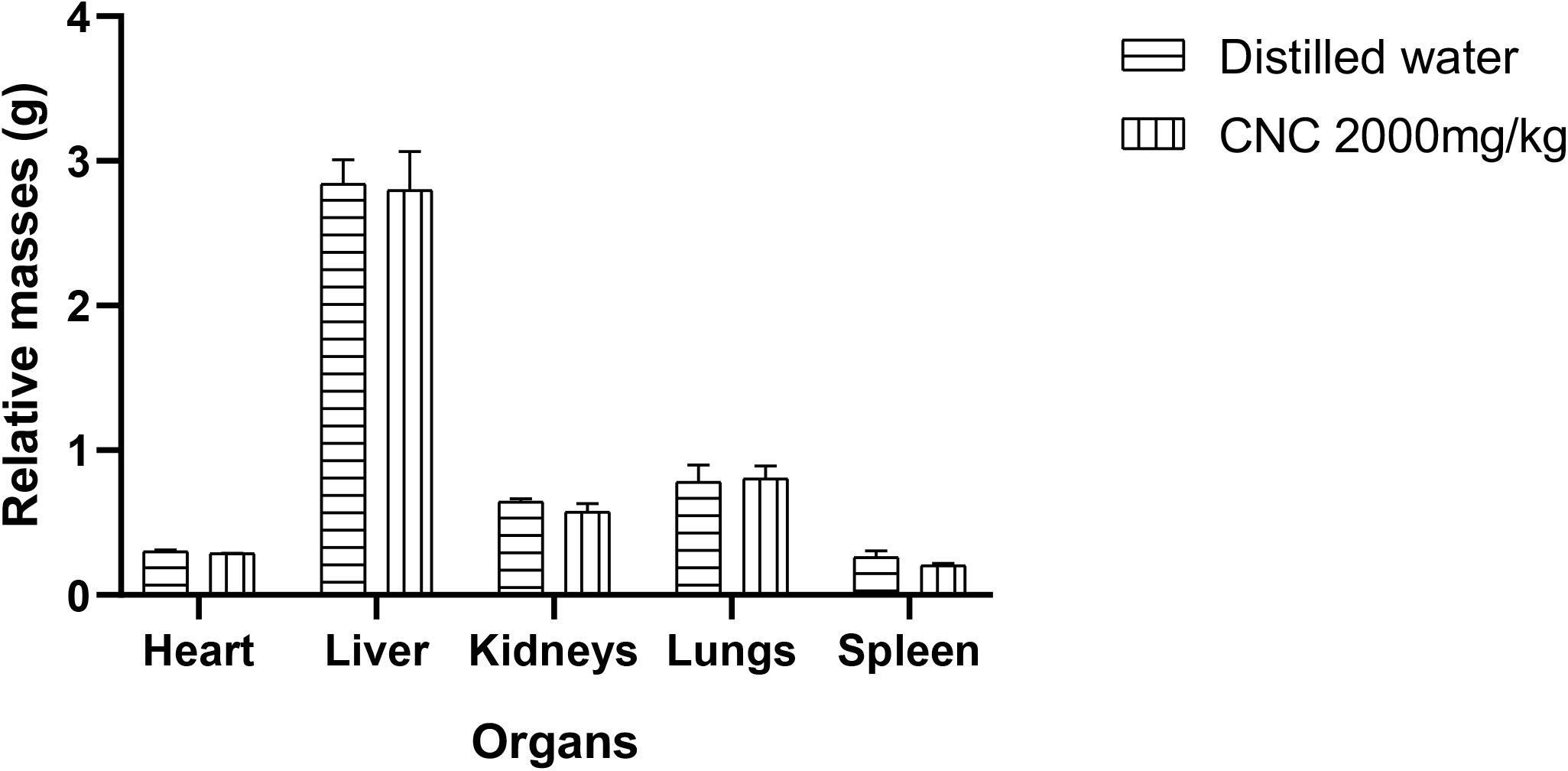
Relative masses of rat organs in test and control groups

### Effect of chitosan nanocapsules on hematological parameters

Immunosuppression was induced in rats by administration of dexamethasone. The animals were treated with different solutions: extract, empty CNC, and CNC with extract. The hematological parameters are presented in Table 5.

**Table 5.** Effect of CNC on the hematology of dexamethasone-induced immunosuppressed rats.

| Hematological parameter | Groups |  |  |  |  |  |  |  |
| --- | --- | --- | --- | --- | --- | --- | --- | --- |
|  | Control batches |  |  | Test batches |  |  |  |  |
|  | Sham control | Negative (H<sub>2</sub>O) | Levamisole | Extract 500mg/kg | Empty CNC 500mg/kg | CNC 100mg/kg | CNC 200mg/kg | CNC 500mg/kg |
| Hematocyte (mm<sup>3</sup>) | 4.1±0.2 | 0.9±0.5 | 0.48±0.05 | 5.35±0.06 | 4.1±0.7 | 5.4±0.3 | 5.3±0.1 | 5.2±0.2 |
| Leucocyte (mm<sup>3</sup>) | 7±1 | 4±2 | 1.47 ± 0.04 | 5±2 | 5.1±0.5 | 10.5±0.2 | 12.0±0.6 | 5±1 |
| Hemoglobin (g/dl) | 10±2 | 7±4 | 22.8 ± 0.5 | 13.3±0.2 | 13.07±0.03 | 13.7±0.6 | 14±1 | 14±1 |
| Hematocrit (%) | 45±2 | 30±5** | 73±2** | 60±3** | 43.4±0.5** | 68.7±0.5** | 64.8±0.5** | 57.5±0.5** |
| MCV (µm<sup>3</sup>) | 88±1 | 78±1 | 14.8 ±0.5** | 88±7** | 91±2 | 107±2** | 107±1** | 89±8* |
| MCH (pg) | 28±1 | 18±2 | 23.1 ± 0.1 | 29±4 | 28±1 | 34±1* | 29±1 | 29±4 |
| MCHC (g/dl) | 32±1 | 26±3 | 129 ± 3 | 34±2 | 33.5±0.3 | 36.9±0.2 | 37.5±0.6 | 37±1 |
| Platelets (mm<sup>3</sup>) | 146.5±0.7 | 131±1 | 1.05 ± 0.04** | 222±54* | 155±2* | 254±32* | 261±42* | 163 ± 24* |
| Lymphocytes (µl) | 1.8±0.5 | 0.4±0.3 | 0.17 ± 0.02 | 1.2±0.5 | 1.4±0.2 | 2±1 | 3.8±0.6 | 19.97 ± 0.00* |
| Middle cells (µl) | 0.50±0.08 | 0.2±0.1 | 0.05±0.02 | 0.44±0.2 | 0.48±0.08 | 0.40±0.21 | 0.4±0.1 | 0.5±0.3 |
| Granulocytes (µl) | 2.8±0.6 | 2±1 | 0.68±0.05 | 2.3±0.4 | 2.40±0.05 | 12.8±0.2 | 12.7±0.6 | 3±1 |

Results are expressed as Mean±SD in respective measurement units; CNC: Chitosan nanocapsules; MCV: Mean corpuscular volume; MCH: Mean corpuscular hemoglobin; MCHC: Mean corpuscular hemoglobin concentration. (\*) p < 0.05; (\*\*) p < 0.01: significant; (\*\*\*) p < 0.001, compared to the negative control.

### Effect of chitosan nanocapsules on delayed-type hypersensitivity response

Cell-mediated immune response caused by chitosan nanocapsules was assessed by delayed-type hypersensitivity (DTH) response. The rats in each group were primed by injection of chicken red blood cells on day 7 of the treatment and challenged on day 14 by subcutaneous injection of chicken red blood cells into the left hind footpad of the rats in each group. The difference in footpad diameter compared to the negative control group showed inhibition of the DTH reaction in rats, revealing the stimulatory effect of chitosan nanocapsules on T cells (Table 6). Levamisole was used as a positive control and showed a significant effect after 24 h. Extract and empty CNC did not show a significant effect, while CNC with extract showed the most significant effect at concentrations of 100 and 500 mg/kg, after 24 h. CNC with extract at 200 mg/kg showed a slightly significant effect from H4 throughout.

**Table 6.** Effect of chitosan nanocapsules on delayed-type hypersensitivity (DTH) response.

| Batch | Difference in foot pad diameter (mm) expressed as Mean±SD |  |  |
| --- | --- | --- | --- |
|  | H4 | H8 | H24 |
| Negative control | 0.04 ± 0.01 | 0.10 ± 0.03 | 0.07 ± 0.01 |
| PC Levamisole 50mg/kg | 0.6 ± 0.5 | 0.4 ± 0.3 | 0.8 ± 0.7** |
| Extract 500 mg/kg | 0.045 ± 0.007 | 0.5 ± 0.4 | 0.3 ± 0.2 |
| Empty CNC 500 mg/kg | 0.10 ± 0.07 | 0.3 ± 0.3 | 0.4 ± 0.3 |
| CNC 100 mg/kg | 0.7 ± 0.6* | 1 ± 1*** | 1.0 ± 0.9*** |
| CNC 200 mg/kg | 0.6 ± 0.6* | 0.7 ± 0.6* | 0.6 ± 0.5* |
| CNC 500 mg/kg | 0.2 ± 0.2 | 0.6 ± 0.5 | 1 ± 1*** |

CNC: Chitosan Nanocapsules ; (\*) p < 0.05; (\*\*) p < 0.01: significant; (\*\*\*) p < 0.001, compared to the negative control.

### Effect of chitosan nanocapsules on humoral immunity

The humoral immune response in rats was assessed by priming the animals via an injection of chicken red blood cells on day 7 of treatment. On day 14, blood was collected, and serum was obtained. The hemagglutination antibody titer was taken as the reciprocal of the highest dilution of the test sera presenting agglutination. Rats who received chitosan nanocapsules at doses of 200 mg/kg and 500 mg/kg showed a significant increase in the antibody titre compared to the negative control group (Figure 5).

**Figure 5.**
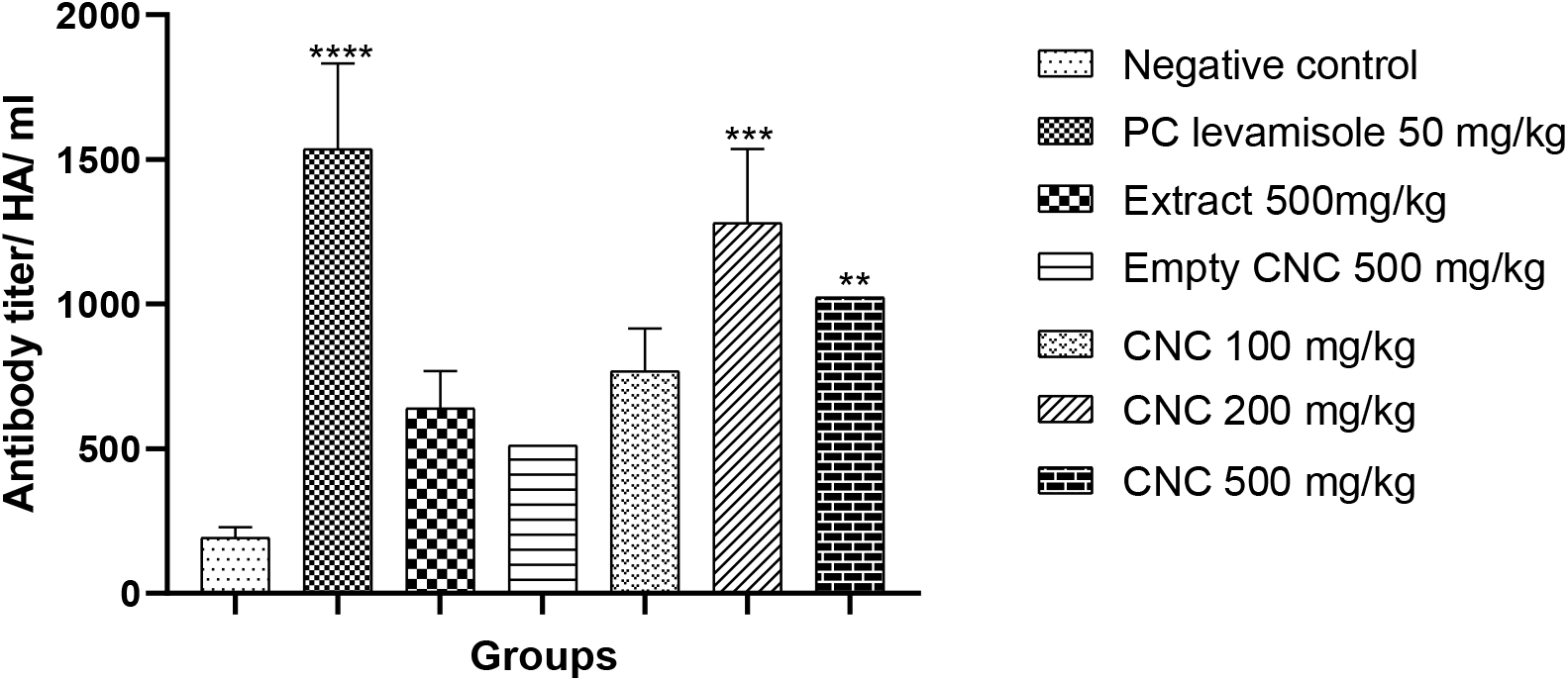
Effect of administration of chitosan nanocapsules on hemagglutination titre. (*) p < 0.05 (**) p < 0.01 (***) p < 0.001 (****) p < 0.0001

## Discussion

The synthesis of chitosan nanocapsules required the extraction using a methanol/dichloromethane solvent system. A yield of 6.11 % was obtained. Adjouzem *et al*. in 2019 obtained an extraction yield of 5.40 % after a single 72 h maceration [15]. The observed variations in extraction yields can be attributed to several factors. The harvest period, which occurred in January in Mbalmayo (Center region) as opposed to March in Loum (Littoral region). The inclusion of dichloromethane enhances the penetration of methanol through the walls of plant cells, thereby improving the extraction of secondary metabolites. Methanol and dichloromethane solvent system, which exhibits affinity for both polar and nonpolar secondary metabolites afford maximum extraction.

Phytochemical screening revealed the presence of polyphenols, namely flavonoids and tannins, as well as alkaloids, steroids, saponins and triterpenoids, as observed by Adjouzem *et al*., in 2019 [15]. Added to these coumarins, reducing sugars and anthraquinones were present. The difference in this phytochemical composition can also be explained by the difference in the harvest period and the different extraction solvent, dichloromethane in this study. The presence of secondary metabolites such as flavonoids, tannins, saponins, and alkaloids explains the immunomodulatory activity observed with the test substances in this study [16]. In view of these results, our extract is rich in secondary metabolic diversity, as observed in other species of the genus *Alstonia*.

The synthesis of chitosan nanocapsules with methanol/dichloromethane extract was carried out by ionic gelation. The efficiency of secondary metabolite encapsulation was obtained by evaluating the degree of encapsulation of total phenols. During ionic gelation, the positively charged chitosan molecules interact electrostatically with the negatively charged tripolyphosphate solution, and a spontaneous electrostatic force of attraction and aggregation follow under the effect of significant rotatory movements. During this process, the secondary metabolites present are encapsulated in a homogeneous manner. The encapsulation efficiency obtained was 69.34%, which depends on factors such as the concentration of the chitosan, of tripolyphosphate solution and the extract, the dispersion rate, and the degree of deacetylation of chitosan. This result corroborates that of Galih Pratiwi *et al*. 2019 [17].

The functional groups present at the surface of the nanocapsules were highlighted by infrared spectroscopy by mechanisms that involve vibrations of elongation and/or deformation of the chemical bonds. The different functional groups observed are consistent with the primary structure of chitosan. As a basic principle, all the functional groups found in the methanol/dichloromethane extract and in chitosan are found in CNC. Practically, because C-O and C-N in chitosan on entrapment generated a new pic characteristic of a new compound which is CNC. This finding is in phase with that of Tchangou *et al*. 2020 [18]. The encapsulated porous material shows a branch of chitosan similar to those described by Gradinaru et al. or Su and colleagues [19,20].

The results obtained for the evaluation of the toxicity of chitosan nanocapsules containing *Alstonia boonei* methanol/dichloromethane extracts reveal low degree of toxicity. The clinical examination and monitoring of rat testicular weight following CNC administration were normal compared to the negative control. Furthermore, the analysis of the relative organ masses revealed no significant differences between the CNC-receiving groups and the control group. Based on these findings, the tested samples can be considered acutely nontoxic at doses below 2000 mg/kg, in accordance with OECD guidelines, which classify substances with an LD_50_ greater than 2000 mg/kg as non-toxic to humans in a single administration. These results are consistent with those reported by Tchangou *et al*. in 2020 [18].

Recently, it has been reported that many natural products have immunomodulatory properties and generally act by stimulating nonspecific and specific immunity. Some of these natural products stimulate both humoral and cell-mediated immunity, while others activate only the cellular components of the immune system. The immune system is the vital defense against non-infectious and infectious diseases. A strong immune system comprises elements that are in balance with each other; if this balance is disturbed, our immune system will not be able to protect the body against harmful invaders [21, 22]. Immunomodulation using natural products can substitute conventional immunotherapy for a range of diseases, especially when the host defense mechanism must be activated under impaired immune response conditions. There are several diseases in which immunostimulant drugs are needed to overcome drug-induced immunosuppression or environmental factors. Drugs that can enhance the immune system to combat the immunosuppressive consequences caused by stress, chronic diseases, and conditions caused by impaired immune responsiveness. Recently, natural products have been commonly used as immunostimulatory agents [23]. Although the natural product has been investigated for various pharmacological activities, the immunostimulatory potential of chitosan nanocapsules remains unknown. In this study result, we showed that chitosan nanocapsules containing A. boonei plant extract had more immunomodulatory activity in experimental models of cell and humoral immunity. The study was carried out using three different methods, each of which provided information about the effect on different components of the immune system. The results of the present study indicate that CNC is a potent immunomodulator, affecting both specific and nonspecific immune mechanisms [24]. The administration of chitosan nanocapsules containing methanol/dichloromethane extract, significantly increased lymphocyte levels, mean corpuscular hemoglobin count, mean corpuscular volume, hematocrit, and platelets count and restored immune deficiency. The results of the present study indicate that chitosan nanocapsules containing *Alstonia boonei* extracts can stimulate bone marrow activity. The bone marrow being the organ most affected during any immunosuppressive therapy [24].

Administration of *Alstonia boonei* methanol/dichloromethane crude extract at 500 mg/kg increased platelets, mean globular volume, and hematocrit, though to a lesser extent than *Carissa congesta* roots, which produced a more significant increase in red and white blood cell counts, as well as in hemoglobin levels at 500 mg/kg [25].

Administration of chitosan nanocapsules at doses of 100 mg/kg, 200 mg/kg, and 500 mg/kg significantly increased hemoglobin, hematocrit, mean corpuscular hemoglobin concentration, and platelet counts, which had been reduced with dexamethasone. This suggests that chitosan nanocapsules can stimulate bone marrow activity. These findings are consistent with those of Wardani et al. (2018), who observed increases in all parameters of total blood count only at a dose of 600 mg/kg of CNC [14].

In the present study, chitosan nanocapsules also showed an overall stimulatory effect on immune functions in rats. Stimulatory effects were observed on both humoral and cellular immunity. Cell-mediated immunity (CMI) involves effectors mechanisms carried out by T lymphocytes and their products (lymphokines). CMI responses are critical to defense against infectious microorganisms, infection of foreign grafts, tumor immunity, and delayed-type hypersensitivity reactions [21,26]. Therefore, the increase in DTH reaction in rats in response to the T cell-dependent antigen due to the mobilization of immune cells at the site of injection brought about by surface antigens of chicken hematocytes, showed the stimulatory effect of chitosan nanocapsules on T cells. In the DTH test, the chitosan nanocapsules showed an increase response in all doses at different time intervals, but this increase was significant only in CNC containing *A. boonei*, but was not significant in groups treated with empty CNC and crude extract. In a similar experiment, *Carissa congesta* roots at 500 mg/kg produced a significant increase in foot pad diameter, a measure of DTH test, which is not in phase with our findings compared to *A. boonei* at 500 mg/kg [14]. These results are like those of Wardani *et al*. in 2018 [14], who obtained a significant increase in foot pad diameter only at doses of 300 mg/kg and 600 mg/kg of CNC. This activity could be due to the synergistic effect between CNC and the extract, amplifying the humoral response by stimulating the macrophages and subsets of B lymphocytes involved in antibody synthesis. The mechanism behind this elevated DTH could be due to sensitized T lymphocytes. When challenged by antigens, they are converted to lymphoblasts and secrete a variety of molecules, including pro-inflammatory lymphokines, affecting more scavenger cells at the site of reaction. An increase in the DTH response indicates a stimulatory effect of CNC that has occurred on the lymphocytes and accessory cell types required for the expression of this reaction [27].

Indirect hemagglutination test was performed to confirm the effect of chitosan nanocapsules containing extract on the humoral immune system. It is composed of B cell with antigens that subsequently proliferate and differentiating into antibodies producing cells, as chicken hematocytes injected into rats having on it surface antigens, stimulated the production and multiplication of specific antibodies.

CNC containing *A. boonei* at 200 mg/kg and 500 mg/kg only, produced a significant increase in antibody titer. In a similar experiment, *Carissa congesta* roots at 500 mg/kg produced a significant increase in antibody titre that is not in phase with our findings compared to *A. boonei* at 500 mg/kg [26]. These results are similar to those of Wardani *et al*., in 2018 [14], who obtained a significant increase in antibody titer only at doses of 300 mg/kg and 600 mg/kg of CNC. CNC at 100mg/kg, empty CNC and crude plant extract were not significant, significance was proven by an increase in antibody titre in rats indicating enhanced responsiveness of B lymphocytes involved in antibody synthesis. This points out the synergistic relationship between CNC and the extract. High values of hemagglutinating antibody titre of the chitosan nanocapsules showed that immunostimulation was achieved through humoral immunity. B lymphocytes and plasma cells function in the humoral immunity component of the adaptive immune system by secreting antibodies such as IgG and IgM, which are the major immunoglobulins that are involved in the complement activation, opsonization, and neutralization of foreign bodies [28].

## CONCLUSION

The synthesis and characterization of chitosan nanocapsules incorporating *Alstonia boonei* bark extract, along with the evaluation of their immunomodulatory activity and acute oral toxicity, in Wistar rats, were presented. Characterization analyzes determined a nanocapusule material with encapsulation rates of 69.34% of the methanol/dichloromethane extract. The chitosan-encapsulated metabolic products exhibited stronger dose-dependent immunomodulatory activity compared to the crude extract and empty chitosan nanocapsules. The nanoderivative showed no adverse effects in acute oral toxicity tests. This study proposes a novel immunomodulatory therapeutic approach that involves encapsulating metabolites in chitosan, leveraging our rich and diverse flora to combat immune diseases.

