## Supplement material 1 for "Chitosan nanocapsules with *Alstonia boonei* extract modulate the immune system in Wistar rats"

Powder X-ray Diffraction analysis (PXRD)

The PXRD of chitosan presents two signals at positions 2 theta 12.9 and 18.6 compared to pure chitosan (9.7 and 20.3°) that represent the diffractions from the (020) and (110) planes of the crystalline lattice (Figure1) [1]. A loss of chitosan crystallinity is observed upon capsule formation. Previous reports on chitosan nanocomposites describe broad peaks that can be shifted together with decrease in crystallinity [2,3]. The PXRD showed a pattern that confirmed the presence of chitosan polymer in the nanocapsule matrix. PXRD confirmed the formation of an organic material composed of C, O and S [2,3].

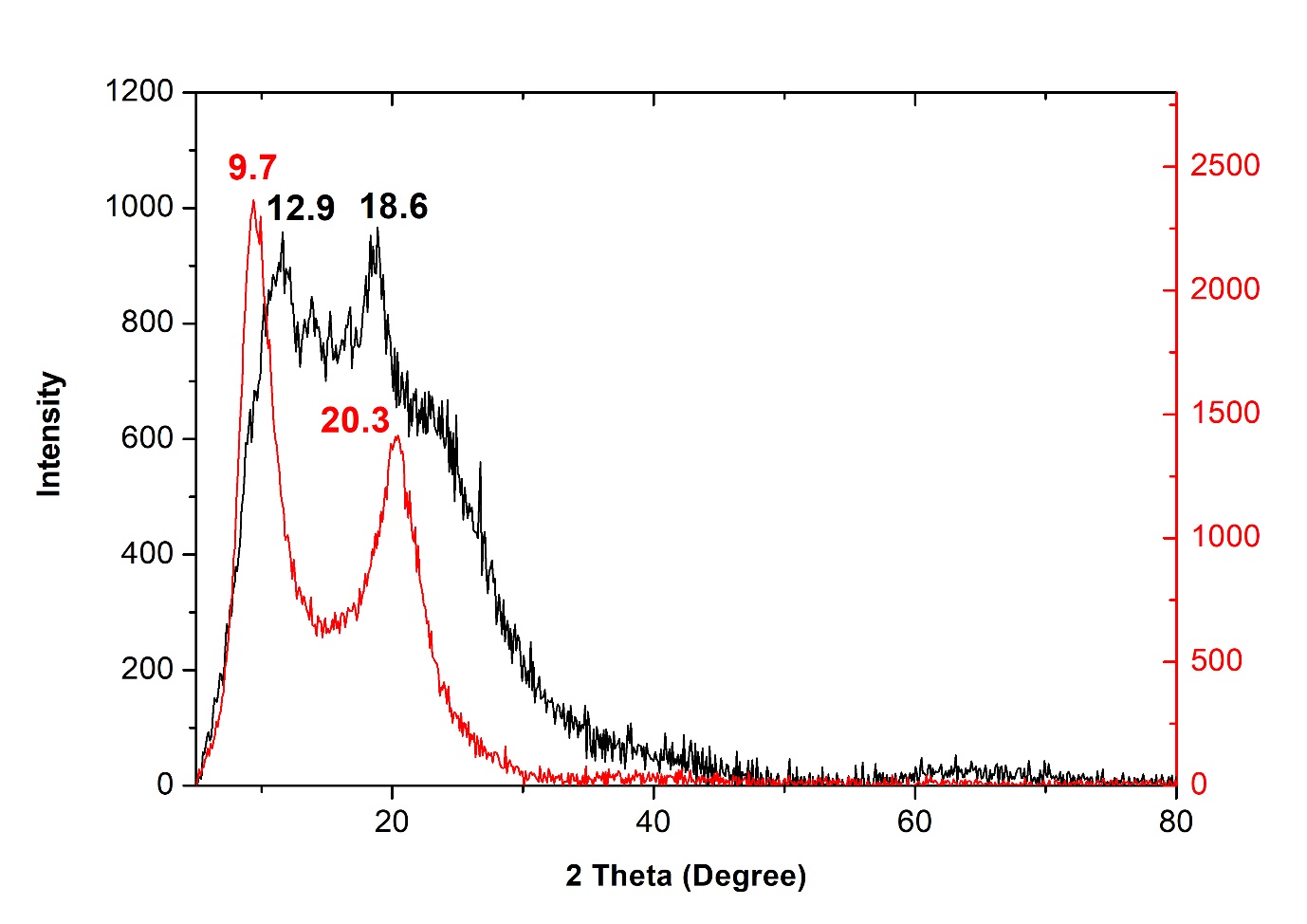

Figure 3. PXRD pattern of Chitosan-*A. boonei* nanocapsules (black) and neat chitosan (red)
